## Supplementary figures for "CHOmics: a web-based tool for multi-omics data analysis and interactive visualization in CHO cell lines"

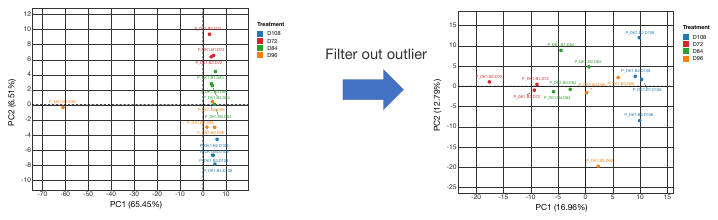

(A) (B)

Figure S1. In (A), PCA analysis on proteomics data shows that one sample at 96 hr is outlier. In (B), the samples are clustered mainly by treatment (i.e., time points) after filtering out the outlier.

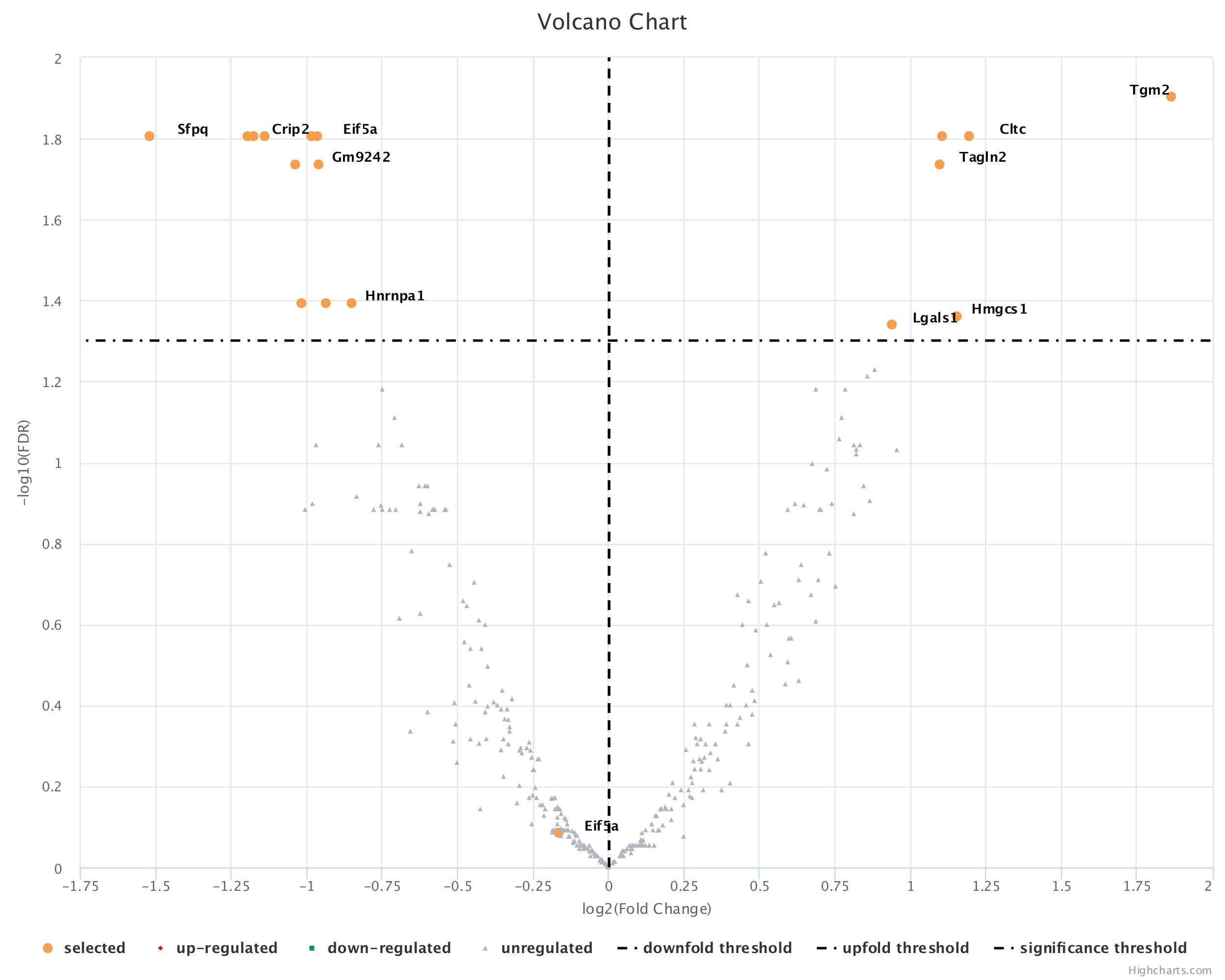

Figure S2. Volcano plot shows the top differential expressed proteins from proteomics data between 108 hr and 72 hr.
